# Inferences about phenological shifts in an Arctic community vary with time-windows

**DOI:** 10.64898/2026.04.13.718090

**Authors:** Patricia Kaye T. Dumandan, Jarno Vanhatalo, Niels Martin Schmidt, Tomas Roslin

**Affiliations:** Department of Ecology, Swedish University of Agricultural Sciences, Uppsala, Sweden; Research Center for Ecological Change, Department of Organismal and Evolutionary Biology, Faculty of Biological and Environmental Sciences, University of Helsinki, Helsinki, Finland; Department of Mathematics and Statistics, Faculty of Science, University of Helsinki, Helsinki, Finland; Department of Ecoscience and Arctic Research Cluster, Aarhus University, Roskilde, Denmark

**Keywords:** Arctic, arthropods, long-term data, plants, phenology, time window, trends

## Abstract

Long-term monitoring data have enabled detection of phenological change, yet it remains poorly understood how its temporal dimensions— duration and choice of start and end years— influence the inferences drawn. To examine which phenological signals emerge at different temporal scales, we analyzed the longest continuous dataset on high-arctic plant and arthropod phenology, collected from 1996 to 2024 in the Zackenberg valley, northeast Greenland. These data have been used to suggest both rapid advancement of spring in the High Arctic (2007) and little directional change but decadal regime shifts (2023). To reconcile these differing conclusions, we quantified how trend estimates varied across moving time-windows and determined the minimum time-series length required to achieve a high probability of agreement with long-term trends. We find that while trend directionality shifts with temporal windows, confidence in trend estimates increases with time-series length. Using the full time-series, we show dampened signals of warming trends, with annual increases in spring and summer air temperatures by 0.04 [-0.05, 0.13] and 0.05 [-0.01, 0.11] °C per year, respectively, alongside a 0.82% [-1.85, 0.19] decline in spring snow cover. We also see modest advancement in the seasonal activity of most arthropod taxa (by ∼0.1 days/year), whereas flowering phenology shows no consistent directional change. Shorter time-series revealed cyclical patterns in abiotic drivers yet variable biotic responses, indicating that a single pattern of “climate change” will translate into varied responses within communities. Finally, almost two decades of data were needed to reliably capture long-term trends. Ecologically, these suggest that 1) phenological shifts in the High Arctic are more moderate than early assessments implied and 2) reflect a dynamic balance between species’ life histories and ongoing climate variability. These may alter interaction potentials within communities, with consequences for ecosystem functioning.

## Introduction

Characterizing trends and drivers of phenology – the timing of recurring biological events – has been a focus of global climate change research (B. Inouye et al., 2019; Thackeray et al., 2010; Visser & Both, 2005). Because species use different phenological cues– from resource species timing their growth to favorable conditions, to consumers using environmental cues that predict resource availability– such studies are key to understanding the ecological impacts of climate change (D. W. Inouye, 2022; Piao et al., 2019). To date, several lines of evidence highlight the role of multiple and interacting drivers – such as temperature, precipitation, photoperiod, and biotic interactions – which also operate across various spatiotemporal scales in shaping phenological patterns (Piao et al., 2019; Thackeray et al., 2010; Wadgymar et al., 2018). For instance, we may expect that drivers that act at broad spatial and long temporal scales (e.g., regional climate patterns) will influence longer-term trends in phenology, whereas drivers that operate at finer scales (e.g., microclimate) may be better predictors of short-term fluctuations in phenology (Park et al., 2021). While there is conceptual support for the need to recognize this scale-dependence, most phenological studies only focus on long-term trends, ignoring the potential of shorter-term trends to reflect meaningful ecological responses.

While estimating long-term trends is undoubtedly important for characterizing complex species’ dynamics, they can however mask substantial variation in phenological responses and limit detection of changing cue-phenology relationships (Iler et al., 2013a). Thus, assessing patterns of shorter-term trends can be useful for capturing nuances around phenological responses to environmental change. Observing systematic variation in shorter-term trends across different time windows may signal change in a species’ sensitivity to particular environmental drivers or changing dynamics of those drivers themselves.

Exploring the incremental change of natural systems through estimation of shorter-term trends is useful not only for identifying temporal patterns of ecological change, but also in addressing practical questions regarding the data demands for effective management. By resampling existing data and estimating trends at different temporal windows, we can gain insights on the value of additional years of monitoring. This is an important consideration for program managers and funders, because collecting information beyond what is necessary could result in delayed action or missed opportunities to act (McDonald-Madden et al., 2010). Thus, the joint characterization of long- and shorter-term trends can be useful for understanding the magnitude and variability of phenological plasticity and the monitoring effort needed to detect it reliably.

The Arctic is an ideal system to explore patterns of phenological plasticity, by estimating long- and shorter-term trends, because of its highly variable and rapidly changing environments (Schmidt, Reneerkens, et al., 2019). This is especially relevant given the rate at which changes in this region are occurring (Rantanen et al., 2022). With a more variable arctic climate (Hartmuth et al., 2023; Kim & An, 2024), the direction and strength of phenological change are unlikely to be sustained through time with fitness consequences shifting across years. This in turn, can make it challenging to predict if and which directional selection is more beneficial.

The long-term record of community phenology in the Zackenberg valley in northeast Greenland offers a case in point. Collected every year since 1996, these data provide the longest, standardised season-wide time-series available on the phenology of high-arctic plants and arthropods. Drawing on different subsets, individual studies have suggested either rapid advancement of spring in the High Arctic (Høye et al., 2007), little directional change but decadal regime shifts (Schmidt et al., 2023) or even sudden breakpoints in the trajectories of individual components (Høye et al., 2021). This begs the question of how the very same dataset can yield so different estimates of change – and whether the different findings can be reconciled without suggesting that one or the other is wrong. In previous works, inferences are drawn based on the full length of the time-series available at the time of the analysis (Høye et al., 2013; Schmidt et al., 2016; Schmidt et al., 2023) and decadal subsets of the data (Schmidt et al., 2023). Moreover, information on the temporal distribution of biotic activity within years has been compressed into simple modes ((Høye et al., 2013; Schmidt et al., 2016); (Høye et al., 2013; Schmidt et al., 2016)23), ignoring potential shifts in the shape of the distribution. These approaches can miss signals that are detectable when resolving finer patterns or using dynamic temporal bounds, and provides us with a limited ability to assess how phenological responses vary at ecologically relevant temporal windows (see (Iler et al., 2013a). Exposing these temporal variations and the underlying mechanisms is also essential for predicting future ecosystem dynamics, as earlier studies have suggested that decreasing overlap between resource and consumer species may be detrimental for either or both parties (Redr et al., 2025; Schmidt et al., 2016) or to arthropod consumers (Høye et al., 2013; Schmidt et al., 2016).

In this paper, we use the Zackenberg monitoring dataset as a case study for extracting more information from phenological time-series, and for reconciling differing findings from the same dataset. Using 29 years of phenology data on plants and arthropods monitored in the Zackenberg Valley with a weekly resolution, we assessed how signals of phenological change are manifested at different temporal scales. To determine long-term trends in Arctic phenology, we sequentially fit hierarchical phenology models with rolling origins to all years of species-specific data. From this, we extracted a global trend parameter that describes the average linear change in peak timing across all years. Then, we visualized and quantified trend uncertainty in two ways: first, we assessed how our confidence in trend estimates vary across different time-windows, and second we identified the minimum time-series lengths (TSLs) at which shorter- and long-term trends tend to agree on the direction of the trend. With this approach, we can determine the scope and detectability of phenological shifts within monitoring programs.

## Methods

### Data

To assess long- and shorter-term trends in Arctic phenology, we used data collected through the BioBasis program under the Greenland Ecosystem Monitoring (GEM) program in Zackenberg, NE Greenland (74°28’N; 20°34’W) from 1996-2024 (Schmidt, Hansen, et al., 2019). Data on environmental variables including air temperature and spring snow cover and on plant and arthropod phenology were accessed from the publicly available GEM database (https://data.g-e-m.dk/).

To characterize abiotic conditions of particular relevance to seasonal patterns of plant and arthropod phenology, we focused on air temperature and snow cover (Koltz et al., 2018; Schmidt et al., 2023). We used air temperature data (°C) collected by two proximal climate stations at 200 cm above ground every 30 minutes. The temperature time series data are the average of the temperatures logged by each sensor. The annual estimate of snow cover in the monitoring area is the mean percent coverage below 600 m a.s.l in the valley observed on June 10 every year (i.e. spring snow cover).

To characterize phenology of the resource populations, we used data on the flowering phenology of *Dryas* (Rosaceae), *Salix* (Salicaceae), *Cassiope* (Ericaceae), *Papaver* (Papaveraceae), *Silene* (Caryophyllaceae), and *Saxifraga* (Saxifragaceae). These data are collected through plant censuses conducted weekly in permanent plant species-specific plots (*Dryas* (n=8 plots), *Salix* (n=6 plots), *Cassiope* (n=6 plots), *Papaver* (n=4 plots), *Silene* (n=4 plots), and *Saxifraga* (n=3 plots)). Data are collected from late May or early June to late August or early September, weather-permitting. In general, samples of a total of at least 50 individuals and their phenological stages (i.e., flower buds, flowers and senescent flowers or capsules with exposed seeds) are recorded within each plot section (For *Salix*, 100 individuals for each stage are counted). More details of the sampling protocol are described in the BioBasis manual (Schmidt, Hansen, et al., 2019). To facilitate data analyses, we only used data collected from June to August for both plants and arthropods, and pooled counts for individuals described to be in non-flowering stages (buds, senescent, exposed seed hairs).

To characterize consumer phenology, we used data on the following arthropod families or superfamilies: *Collembola, Ichneumonidae, Acari, Lycosidae, Sciaridae, Phoridae, Nymphalidae, Muscidae, Linyphiidae, Coccoidea, and Chironomidae*. These taxa are monitored through trapping with yellow pitfalls in permanent trapping stations.

Data cleaning and manipulation steps were done following (Schmidt et al., 2023). For arthropods, we selected the four pitfall trapping stations continuously monitored throughout the study period (Art2, Art3, Art5, Art7). To represent the univoltine nature of most arthropods in the Arctic, we conducted further data filtering by excluding plot-year combinations for each taxon if asymmetric phenological curves or multimodal distributions were found using Hartigan’s dip test for unimodality (Gerlich et al., 2025).

### Data Analysis

#### Peak phenology estimation

To estimate peak phenology, we fitted linear mixed effects models to species-specific data. For plants, where phenological stage had been usually scored in three categories (buds, inflorescences and senescent inflorescences), each model had the form:

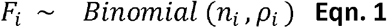

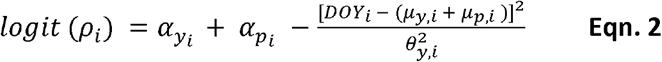

Here *F*_*i*_, the number of flowering individuals out of total individuals observed, is assumed to follow a binomial distribution with *n* (number of trials, e.g., number of flowers and non-flowers) and *ρ* (probability of success, e.g., the probability of flowering individuals; Eqn. 1) parameters. The mean probability of flowering, *ρ*, is modelled as a function of year-level 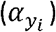 and plot-level intercepts 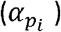, and the alternative formulation of the quadratic term for day of year (*DOY*) in terms of the mean (*µ*; interpreted as peak timing) and width (*θ*; interpreted as a measure of length of the flowering season) parameters. (Eqn. 2). Peak timing was modeled using a non-centered hierarchical parameterization, with standardized latent variables scaled by population-level standard deviations. Plot-level deviations in peak timing were modelled as: 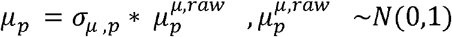, which allowed peak timing to vary among plots. Year-specific peak timing was estimated as a linear function of time (year) with interannual variability: 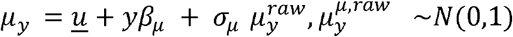 The global trend in peak timing is represented by *β*_*µ*_, and year-level deviations from this trend are captured by year-level random effects. Finally, phenological width was also allowed to vary per year: 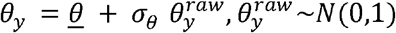

The general model form for arthropods was similar but we assumed the data to follow a negative binomial distribution (Eqn. 3), with mean (*η*_*i*_) and overdispersion (*τ*) parameters, and included an offset term for observation effort (days, *log* (*K*_*i*_) ; Eqn. 4).

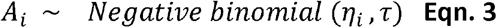

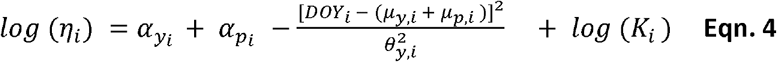

#### Trend estimation

To assess the role of time-series windows (i.e., length and starting year) on signals of phenological change, we iteratively refit the models described above across rolling-origin windows (Fig. 1), and extracted posterior draws of the trend parameter for peak timing (*β*_*µ*_). This produces a distribution of slopes that captures both within-window variability and posterior uncertainty. In doing so, we track how the slope estimates (i.e., temporal trends) and its uncertainty change as the time-series start year moves forward, and the time-series length increases. Window sizes were initially set at five-years, and allowed to grow incrementally by a year at a time.

**Fig. 1.**
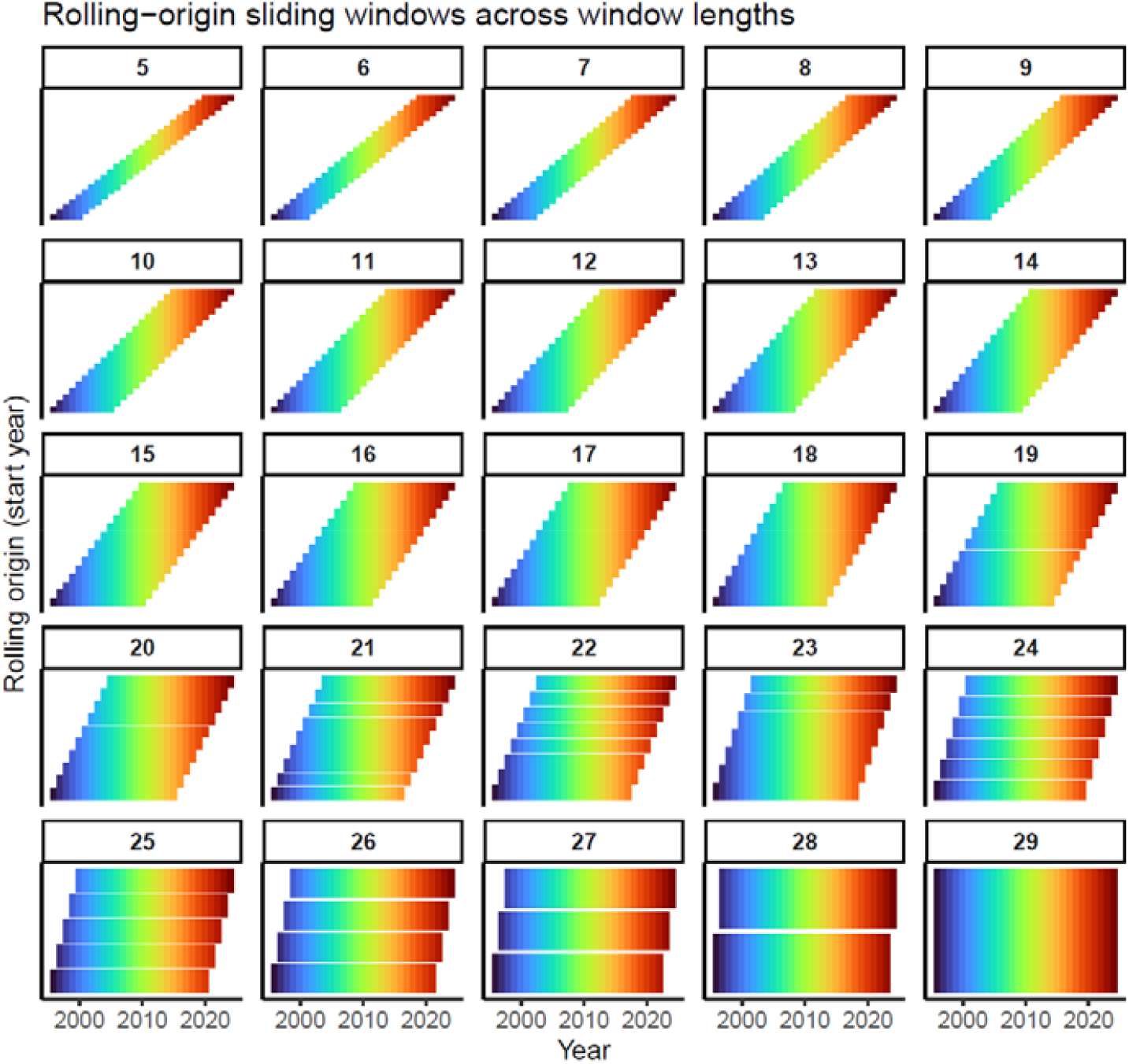
Conceptual summary of the study design and analytical approach. To estimate trends in peak phenology with rolling origins, we used a sliding window analysis (minimum time window was 5 years). Each panel shows rolling windows for a fixed window length. Each horizontal row in each panel represents a rolling-origin window beginning in a specific start year and extending forward for the duration of the window. Colored tiles indicate years included in each window, with blue representing earlier years and red for later years.

#### Output interpretation

To determine how the directionality and strength of the signal of phenological change varies with choice of temporal window, we used the slope estimates from each model at each origin. Specifically, we used the mean slope and the width of its 95% credible interval (i.e., difference between the upper (97.5) and lower (0.025) quantiles) to gain intuition on the magnitude of certainty, and interpreted the inverse of this value as a slope estimate with higher certainty. To assess our confidence in trend estimates, we used the inverse of the 95% credible interval of the trend parameter — with smaller values interpreted as trends with higher certainty. Then, to determine the scope and detectability of phenological shifts within monitoring programs, we identified the minimum time-series lengths (TSLs) at which there is high probability of agreement in directionality between shorter- and long-term (i.e., full time-series) trends. To identify the minimum TSL required to recover the direction of long-term phenological change, we used the posterior draws of slope estimates fitted across multiple time-windows. For each window, we quantified the probability of agreement between the shorter- and long-term trends as the proportion of posterior draws sharing the same sign as the long-term slope. We then aggregated these proportional values across windows of equal TSL, and defined the minimum threshold as the shortest TSL at which the posterior probability of agreement was ≥ 0.95. This approach effectively incorporates posterior uncertainty while accounting for within-window variation.

#### Model implementation and diagnostics

All models were implemented in a Bayesian framework. Parameters inside the quadratic term follow a non-centered hierarchical structure, and along with all global parameters, were given weakly informed, normally distributed priors with a mean of 0 and a variance of 1 or 2. We based parameter estimates from models that were run on four chains of 2,000 iterations, with 500 used in the burn-in period. We visually assessed model convergence and performed posterior predictive checks to evaluate deviations of model-generated data from the observed data. We interfaced to Stan using the ‘rstan’ package ver 2.32.6 (Stan Development Team, 2024). All data curation and analyses steps were done using R ver. 4.3.0 (R Core Team, 2023). Analysis scripts are deposited in a <u>Github repository</u>, which will be deposited in Zenodo upon manuscript publication.

## Results

### Environmental conditions

On average, long-term trends of the three covariates examined in this study suggest warming conditions in the Arctic. Spring and summer air temperatures were increasing by 0.04 [-0.05, 0.13] and 0.05 ℃ [-0.01, 0.11] annually, and spring snow cover contracted at an annual rate of 0.82% [-1.85, 0.19]. These similar warming trends, particularly for summer air temperature and snow cover, were more probable during earlier years of monitoring (late 1990s to mid 2000s, Fig. 2) when inspecting shorter time-series. On the other hand, short term trends of spring air temperature seem to shift direction approximately every five to seven years, with cooling phases being relatively shorter (Fig. 2). While we did not find a relationship between trend certainty and time-series length for any of the abiotic variables, we do note oscillations in the trends of both spring air temperature and snow cover, and more mixed trends in summer air temperature during the more recent period of monitoring (Fig. 2).

**Fig. 2.**
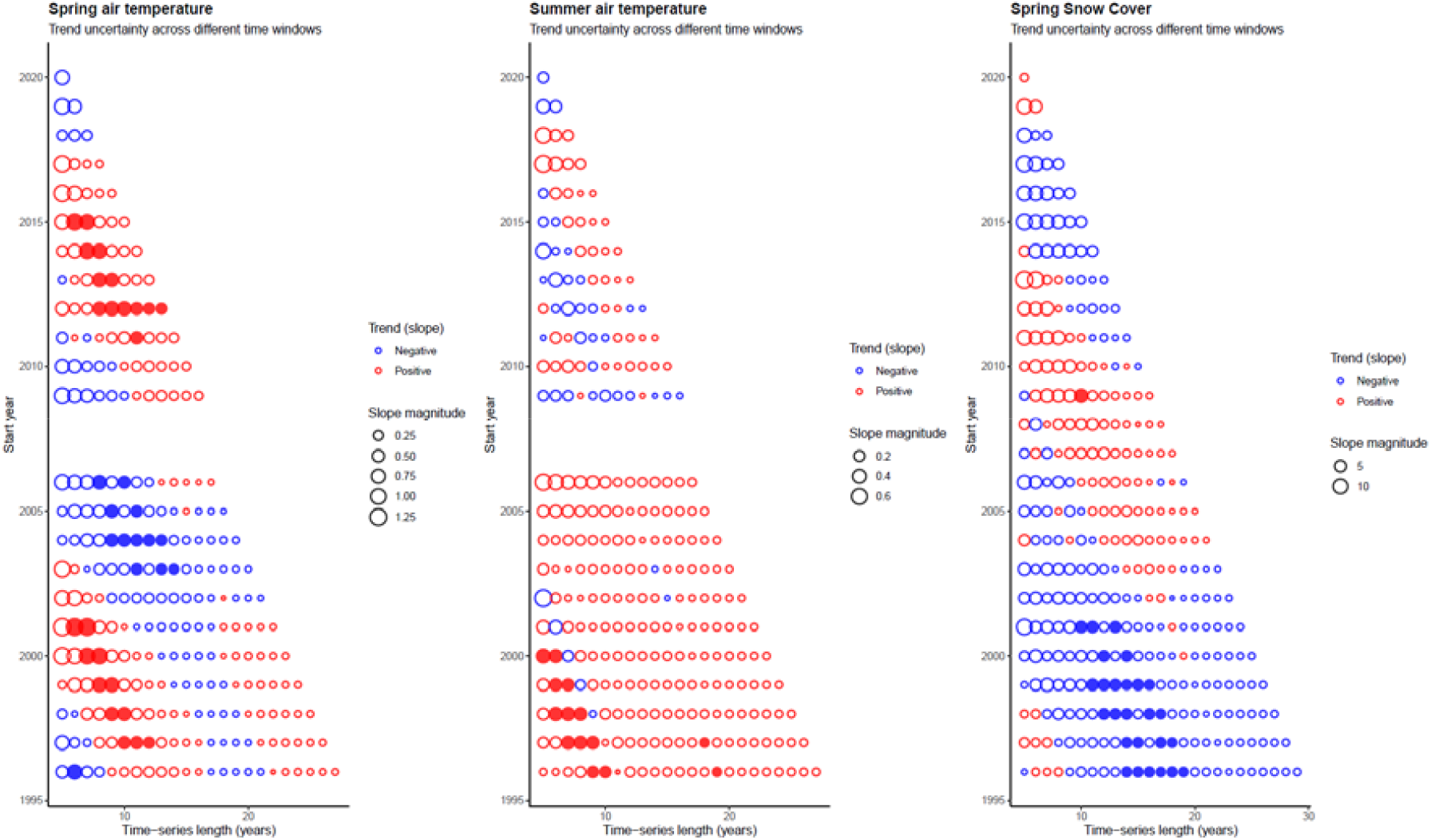
Trends in the key environmental covariates for Arctic plants and arthropods: spring and summer air temperature, and spring snow cover (left to right panel). Each bubble represents a slope estimated from a linear regression as the time-series start year moves forward, and the time-series length increases. The minimum time window was 5 years. Colors and fillings of the bubbles indicate trend directionality and signal strength. Blue bubbles show cooling (or decreasing) trends while red indicates warming (or increasing) trends in air temperature (or snow cover), with filled ones indicating significance. Bubble sizes indicate slope magnitudes, with larger bubbles indicating larger year effects. Gaps indicate absence of a slope from a specific window.

### Plant and arthropod phenology

Long-term trends in peak phenology varied among taxa. In general, signals of phenological change, when considering the full time-series were dampened for both plants and arthropods, although directional change was more consistent among arthropods than among plants. On average, arthropods have advanced their phenology by approximately 0.1 days/year (Fig. 3). Herbivorous arthropods, *Nymphalidae* and *Coccoidea*, advanced their phenology at a rate of 0.23 [-0.44, -0.02] days/year and -0.03 [-0.18, 0.12] days/year, respectively. The aquatic *Chironomidae* also advanced its phenology by -0.07 [-0.18, 0.04] days/year. Among opportunistic predators, most groups except *Linyphiidae* (1.30 [-0.38, 2.32] days/year) advanced their phenology (*Lycosidae* (-0.09 [-0.29, 0.11] days/year) and *Ichneumonidae* (-0.01 [-0.13, 0.11] days/years)). Soil-eating and/or detritivorous arthropods including *Acari* (-0.01 [-0.11, 0.10]), *Collembola* (-0.15 [-0.34, 0.03]), *Muscidae* (-0.09 [-0.22, 0.03]), *Phoridae* (-0.15 [-0.28, -0.03]), and *Sciaridae* (-0.11 [-0.26, 0.05]) have also shown, on average, advancing phenological trends (Fig. 3).

**Fig. 3.**
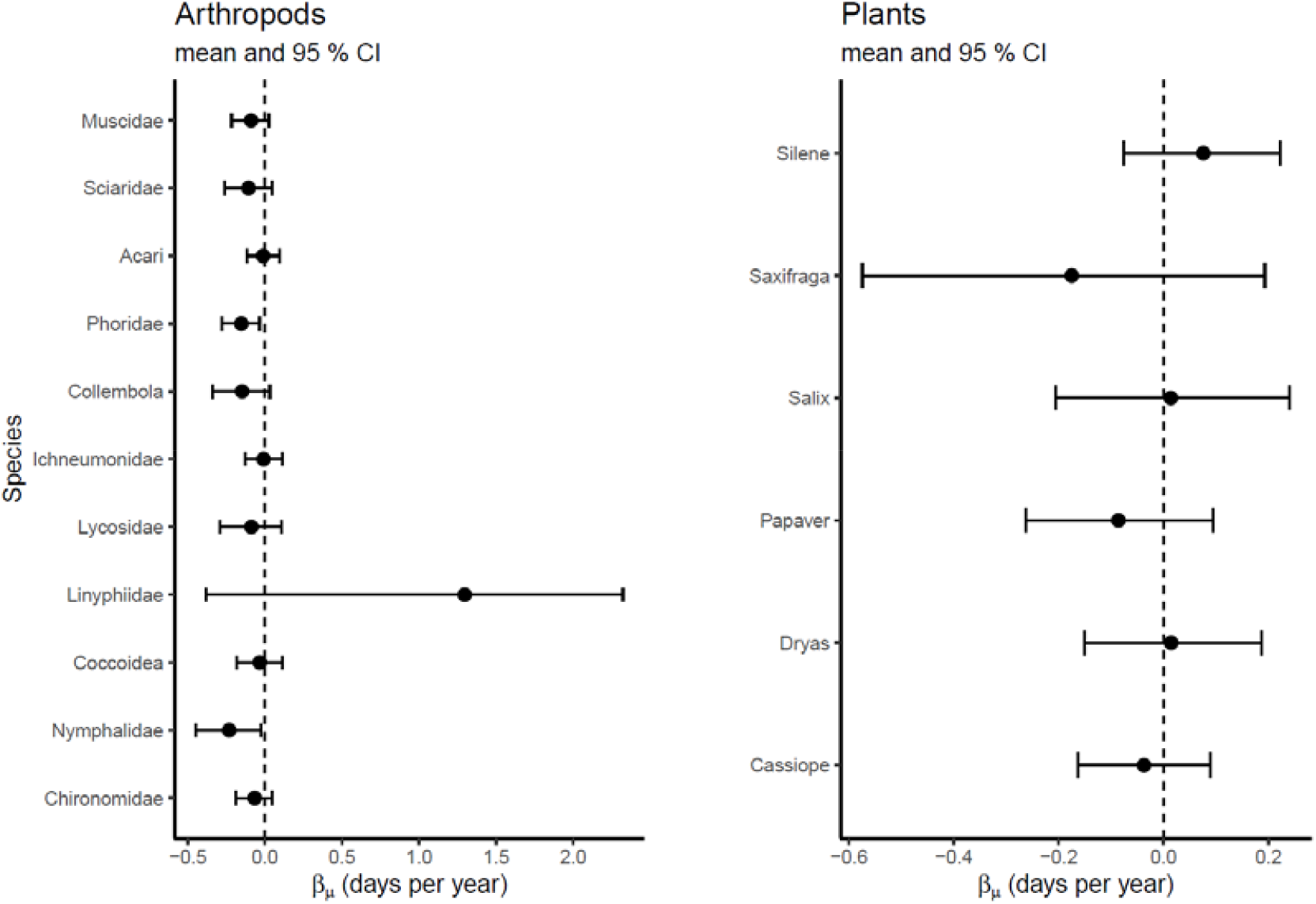
Long-term trends in peak phenology of arthropod and plant taxa. Whisker plots depict the mean and 95% CI of the parameter, which represents the long-term trend for each taxa.

Unlike arthropods, directional selection was less evident among plants. *Cassiope* (-0.04 [-0.16, 0.09]), *Papaver* (-0.09 [-0.26, 0.10]), and *Saxifraga* (-0.18 [-0.57, 0.19]) exhibited advancing trends while *Dryas* (0.15 [-0.15, 0.19]), *Salix* (0.01 [-0.21, 0.24]), and *Silene* (0.08 [-0.07, 0.22]) exhibited delaying trends, on average (Fig. 3).

When examining shorter subsets of these species-specific data, we did not detect a pattern in directional selection across taxa (Fig. 4). However, we found a clear effect of time-series length on trend certainty: longer time-series were associated with increased certainty in slope estimates (Fig. 5). This indicates that while the probability of inferring advancing or delaying trends is variable with shorter time-windows, the reliability of trend estimates increases with time. In both plants and arthropods, the minimum time-series length required to observe agreement between long- and shorter-term trends exceeded 10 years (Fig 5). Among representative taxa, the influence of start year on within-window variability was most pronounced in *Silene*, reflected by the large variation in the 95% CI width of the slopes across different start-years (Figs. 4 and 5).

**Fig. 4.**
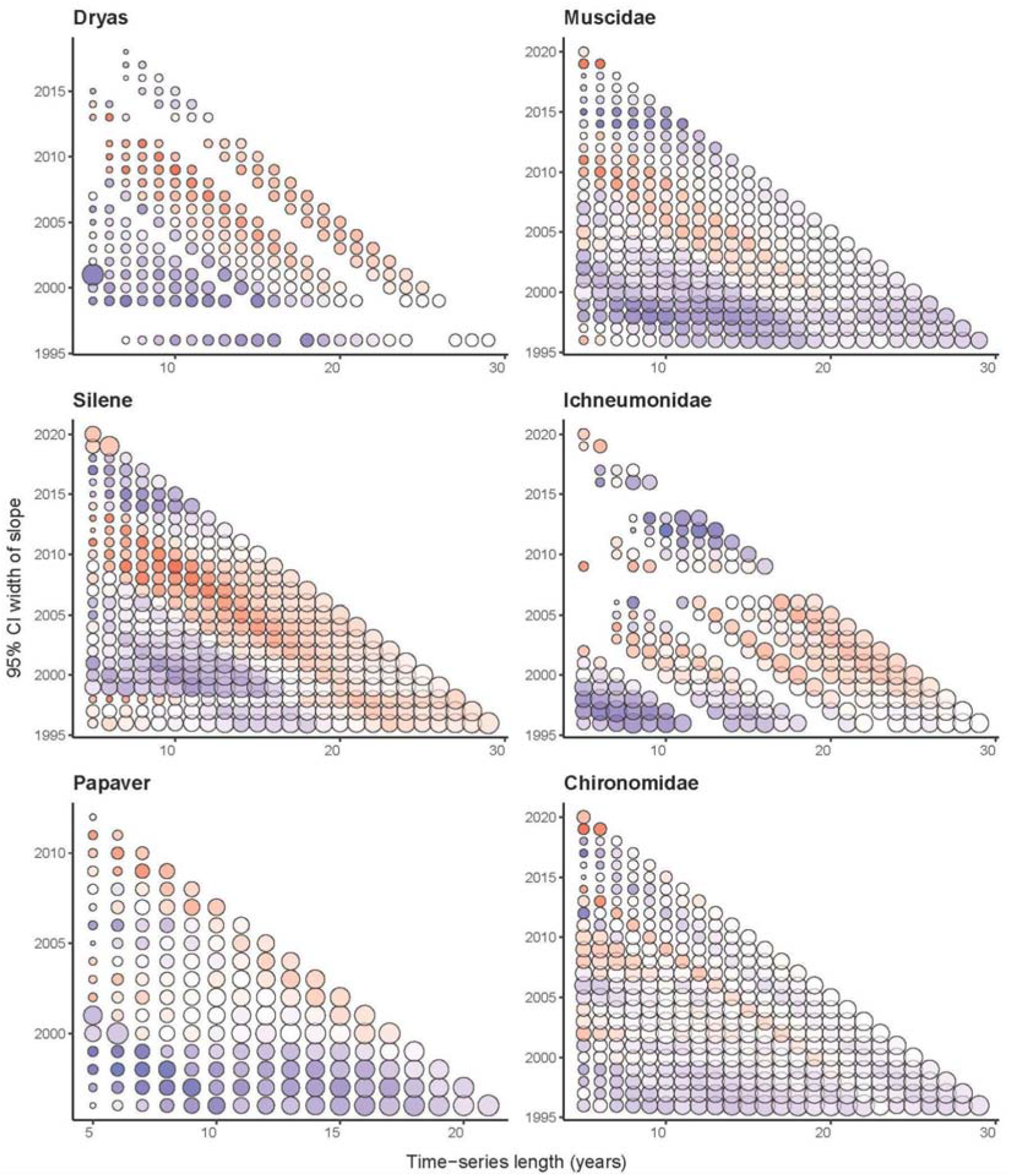
Temporal trends observed at various time-windows for representative plant (left panel) and arthropod (right panel) taxa. The minimum time window was 5 years. Each bubble represents a slope estimated from a sequential model fit. Variation in bubble color and shading indicates directionality and strength of signal of the trend. Darker blue suggests advancing trends while darker red suggests delaying trends. Bubble size indicates the inverse of the 95% credible interval of the slopes, with larger circles representing higher trend certainty. Gaps indicate absence of a slope from a specific window. Representative taxa were selected based on qualitative differences in their apparent phenological patterns to illustrate the range of variation observed across taxa.

**Fig. 5.**
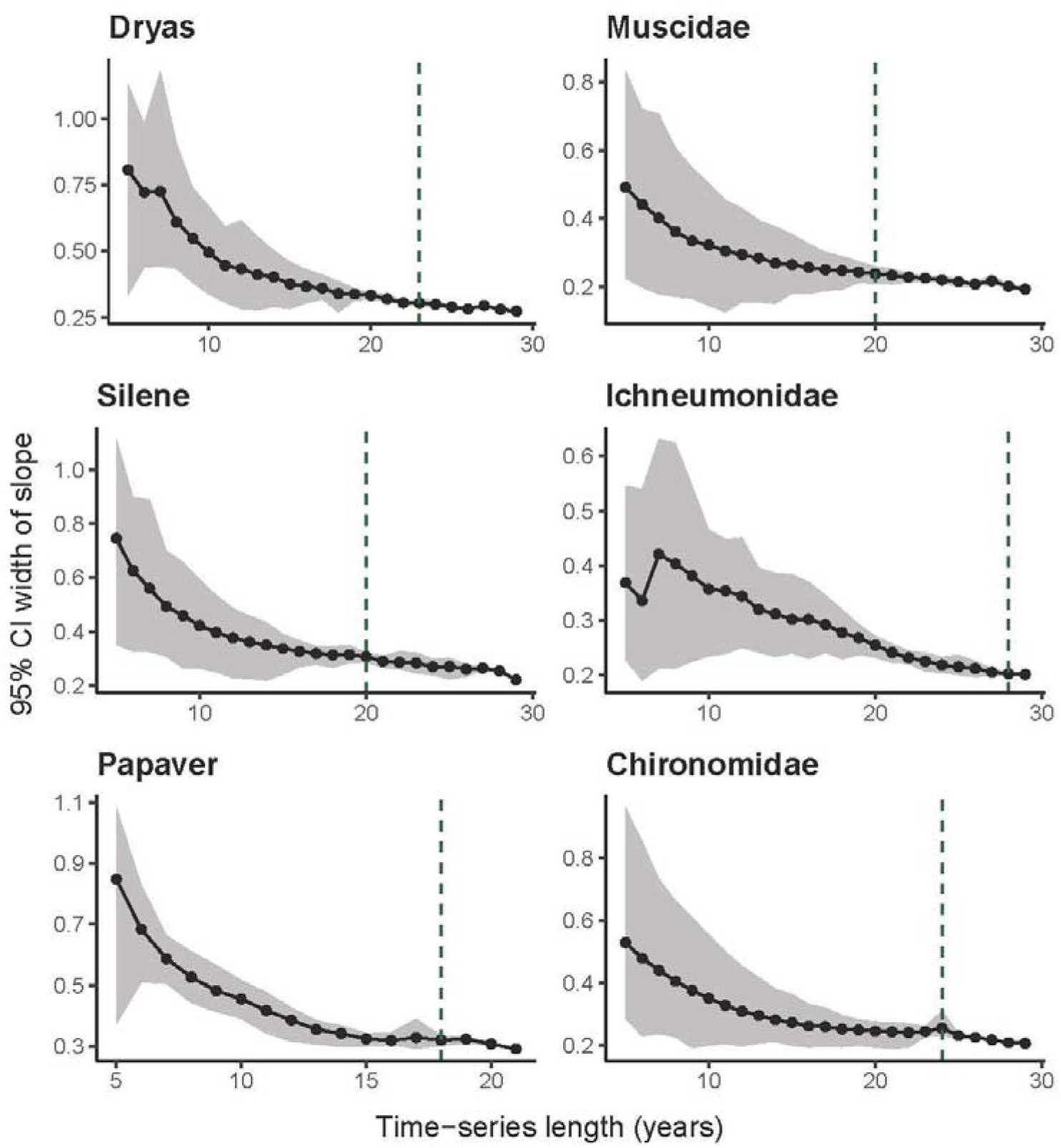
Trend uncertainty over time for representative plant and arthropod taxa. Each point indicates the mean difference between the upper and lower 90% bounds of the slope estimated for a given time-series length, while the gray shading around the line indicates the 95% CI. Dashed lines indicate the minimum time-series length required to recover the direction of long-term phenological change (estimated from full dataset).

## Discussion

Our perceptions of the magnitude and direction of ecological change are strongly shaped by the temporal bounds used to describe it. Shorter vs. longer time-series will be differently reflective of short-vs long-term dynamics in the data (Iler et al., 2013a), and it is mainly when we use time-series with suboptimal lengths to characterise long-term dynamics that inference may be compromised (Høye et al., 2007). Here, we demonstrate that inferences about changes in arctic phenology depend largely on the choice of time-window. When shorter time-windows (< 10 years) are considered, we find higher variation in trend certainty across different start years for most taxa (Fig. 4). We also note group differences in long-term trends. When trends were assessed over long time-scales (i.e., full time-series), plant taxa were less likely to exhibit directional change than arthropods (Fig 3). Nonetheless, shifts in trend directionality is apparent in both plant and arthropod taxa, as well as for abiotic variables (Fig. 2 and 4). This suggests that interannual variability and periodic dynamics can be obscured by long-term trend estimates. Finally, as theoretically expected, we provide empirical evidence that longer time series increase the certainty of trend estimates (Fig. 5).

The current findings provide a possible explanation for why early inferences about rapid advancement of phenology (Høye et al., 2007) have later been replaced with reports of much lower rates of change (Schmidt et al., 2023)). The start and end years used in the earlier analysis (Høye et al., 2007) coincided with a negative phase of the North Atlantic Oscillation (NAO), which is often correlated to warmer temperatures in the region (Folland et al., n.d.; Hanna et al., 2015), and is, in turn, conducive for earlier onset of seasonal activities of many species. Summer NAO have become more variable in the more recent past (Liu et al., 2025), which may have played a part in the dampened signals recovered by more recent studies (Schmidt et al., 2023)), including this one. This observation is fully consistent with other work on the effect of time-series length, and the choice of specific start and end years. An outlying data point that occurs earlier or later in the time-series can exert a disproportionately large impact on estimated linear trends (White, 2019). Indeed, we find multiple examples of how different start years can yield different narratives of phenological change. For example, the decadal trends for *Silene* and *Papaver* estimated earlier in the monitoring period (late 1990s) were different from those estimated in the mid-2000s (2005-2010; Fig. 4). We observed similar contrasts for *Muscidae* and *Chironomidae*. These periods that seem to have contrasting trends may reveal system-specific idiosyncrasies, such as shifts in monitoring protocol, extreme weather events or other transient dynamics. Once sufficient data have been accumulated, analyses such as ours can be used to efficiently explore the sensitivity of trend estimates to particular choices. By examining time-series data across different temporal windows (i.e., start years), we emphasize the importance of long-term datasets not only for capturing slow-unfolding ecological processes, but also shifts in the variability of natural systems (Gauthier et al., 2024; Lindenmayer et al., 2022; Murray & Krebs, 2024).

This is not to say that short-term dynamics are redundant and long-term trends are all that matter because it is quite the opposite. In this study, we show the multitude of patterns emerging at the level of individual taxa (Fig. 4), and in comparisons between trends in abiotic conditions (Fig. 2) and biotic responses (Fig. 4) when shorter time-series are examined. Indeed, these results suggest that species response will be dictated by the match between an organism’s life cycle and the time span of shorter-vs. longer-term climate dynamics. Of the arthropod species studied here, most will live for two to a few years (Elkington, 1971; Schmidt et al., 2006), whereas most of the plants, except *Papaver*, will live for decades or centuries (Böcher et al., 2015; Schmidt et al., 2006). Thus, the fitness and phenology of an arthropod cohort may be dictated by climatic variation at a much shorter scale than its resource species will respond to. One and the same pattern of “climate change” will thus translate into varied responses among community members (Iler et al., 2013a).

Direct associations between species’ responses and environmental conditions are often short-lived (compare Fig. 2 and 4) because many species integrate environmental cues over specific, and sometimes narrow time-windows rather than responding to long-term climatic averages (Watts et al., 2022). This further highlights the value of characterizing trends not only at longer time scales but also at shorter time-windows, because the latter may capture a different aspect of environmental forcing than longer ones. In this study, this is demonstrated by the lack of persistent correspondence between abiotic trends and phenological responses. For example, warmer temperatures and reduced snow cover were not always matched by advances in plant or arthropod phenology (Fig. 2 and 4). We also observed lower variability in summer air temperatures during earlier years of monitoring; however, around the 2010s, both short- and long-term trends exhibited increased variability (Fig. 2). In contrast, we did not detect a comparable periodicity in the phenology of arctic plants and arthropods. Taken together, these may suggest that while climate is a key indicator of community phenology in the Arctic, its importance may be context-dependent. These results highlight the importance of mechanistic models to better understand species’ response to novel and increasingly variable climatic conditions (Railsback et al., 2025).

From a monitoring perspective, our findings also reveal how long time-series data should be to reliably estimate phenological change. Results shown here reveal that the persistence of trend directionality has frequently weakened within less than a decade for many taxa (Fig. 4). This manifests a nonlinear phenological response, which can be obscured with traditional approaches to estimating trends (Høye et al., 2021; Iler et al., 2013b; Rafferty et al., 2020). For example, signals of shifts in trend directionality in representative plant, *Silene*, and arthropod, *Muscidae*, appeared less pronounced after approximately five years of data beginning in 1997. These results demonstrate that both the detectability and stability of phenological signals can vary substantially through time, even within long-term datasets.

Although decay in phenological signals over time has been well-documented (see also below), we add nuance to it and show that while signals can get dampened in long-term trends, uncertainty decreases (Iler et al., 2013a, 2017; Schmidt et al., 2023); Fig. 5). By estimating the minimum time-series length required to achieve a high probability of agreement in trend directionality between shorter- and long-term trends, we suggest that the current recommendation of at least a decade-long monitoring to estimate phenological trends may not be sufficient (Parmesan, 2007; Thackeray et al., 2016). For many taxa, more than 20 years of data were needed to confidently infer the underlying (“true”) trend (Fig. 5). Here, one may naturally argue that the current time series of 29 years may still be too short to reveal the true trend – but the narrow confidence intervals achieved (Fig. 5) suggest that we can at least be reasonably confident about our current estimate.

Overall, these results underline how inferences about phenological trajectories are influenced by time-windows. As such, they are important considerations for the interpretation of any monitoring data. At the same time, they raise the obvious question of *when one might stop monitoring*. This question seems especially relevant for program managers, given that the maintenance of long-term monitoring sites requires balancing costs against scientific insights gained (Bennett et al., 2018). The monitoring data collected in the Zackenberg Valley for almost three decades has been instrumental for revealing Arctic change, and has motivated several lines of inquiry that strengthen our ability to predict biodiversity change under different climate change scenarios (Meltofte et al., 2021; Roslin et al., 2013; Schmidt et al., 2017). At the same time, the current results might be read as suggesting that a “surplus” in data being collected at Zackenberg – given that we have already exceeded the minimum time-series length required for shorter – and long-term trends to match. This *might* be true *if* climate change had stopped – but it clearly has not, and future conditions may differ substantially from current ones (Liu et al., 2025; Schmidt et al., 2023). Our results should therefore be interpreted as data length is sufficient for now – by revealing the true rate of change so *far*. However, future environmental changes may result in a complete revamp of this “true” long-term trend. Thus, we continue to emphasize the importance of long-term monitoring programs as a pivotal tool to map and understand ecosystem dynamics in a rapidly-changing world.

## Author Contributions

**Patricia Dumandan:** conceptualization, formal analysis, writing-original draft. **Jarno Vanhatalo: formal analysis**, writing- review and editing. **Niels Martin Schmidt:** data curation, writing-review and editing. **Tomas Roslin:** conceptualization, funding acquisition, writing- review and editing.

## Acknowledgments

We thank the past, present, and future Greenland Ecosystem Monitoring and in particular BioBasis staff at Zackenberg Research Station for collecting data over the years. Funding awarded to TR by the Swedish Research Council (2023-05118) supported this work. PD received training in Bayesian modeling supported by the National Science Foundation (2042028), which has enhanced the work reported here. JV was funded by the ERC Consolidator Grant (BEFPREDICT, 101087409).

## Data Availability Statement

Data on environmental variables including air temperature (https://doi.org/10.17897/G5WS-0W04) and spring snow cover (https://doi.org/10.17897/M7Y2-TK96), and on plant (https://doi.org/10.17897/X0MY-K003, https://doi.org/10.17897/JSQ7-6355, https://doi.org/10.17897/NK32-H804, https://doi.org/10.17897/NS7W-JT18, https://doi.org/10.17897/YXH1-ZB25, https://doi.org/10.17897/6GVG-QH42) and arthropod (https://doi.org/10.17897/V285-Z265) phenology were accessed in the publicly available GEM database (https://data.g-e-m.dk/). Analysis scripts are currently deposited in a Github repository (https://anonymous.4open.science/r/ZackPhen/README.md), but will be deposited in Zenodo upon manuscript acceptance.

