## Supplementary Materials for "Inferences about phenological shifts in an Arctic community vary with time-windows"

**Running title:** Phenological change across time windows

**Supplementary Figures**

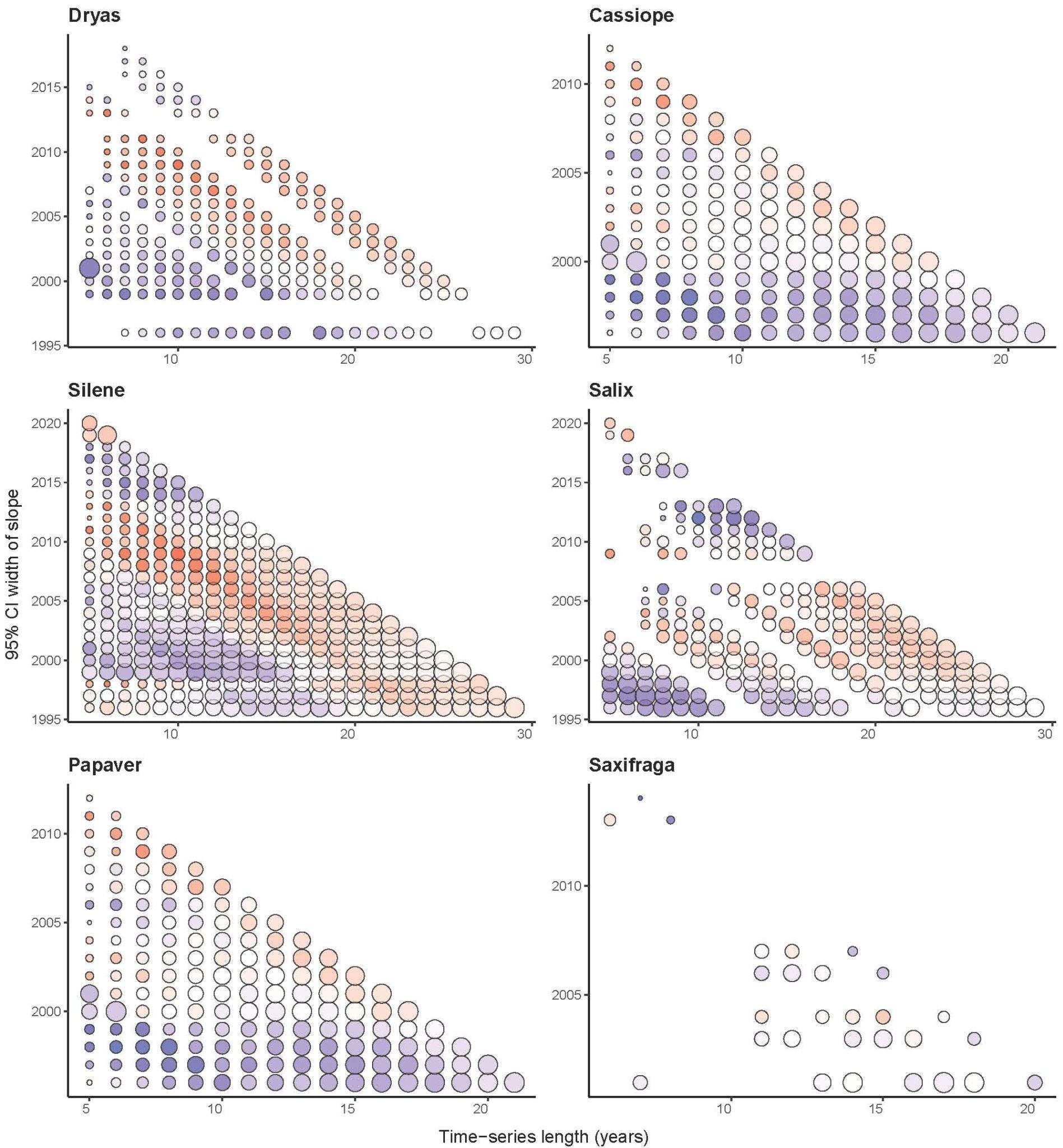

Supplementary Fig. 1. Temporal trends observed at various time-windows of all plant taxa. The minimum time window was 5 years. Each bubble represents a slope estimated from a sequential model fit. Variation in bubble color and shading indicates directionality and strength of signal of the trend. Darker blue suggests advancing trends while darker red suggests delaying trends. Bubble size indicates the inverse of the 95% credible interval of the slopes, with larger circles representing higher trend certainty. Gaps indicate absence of a slope from a specific window. Representative taxa were selected based on qualitative differences in their apparent phenological patterns to illustrate the range of variation observed across taxa.

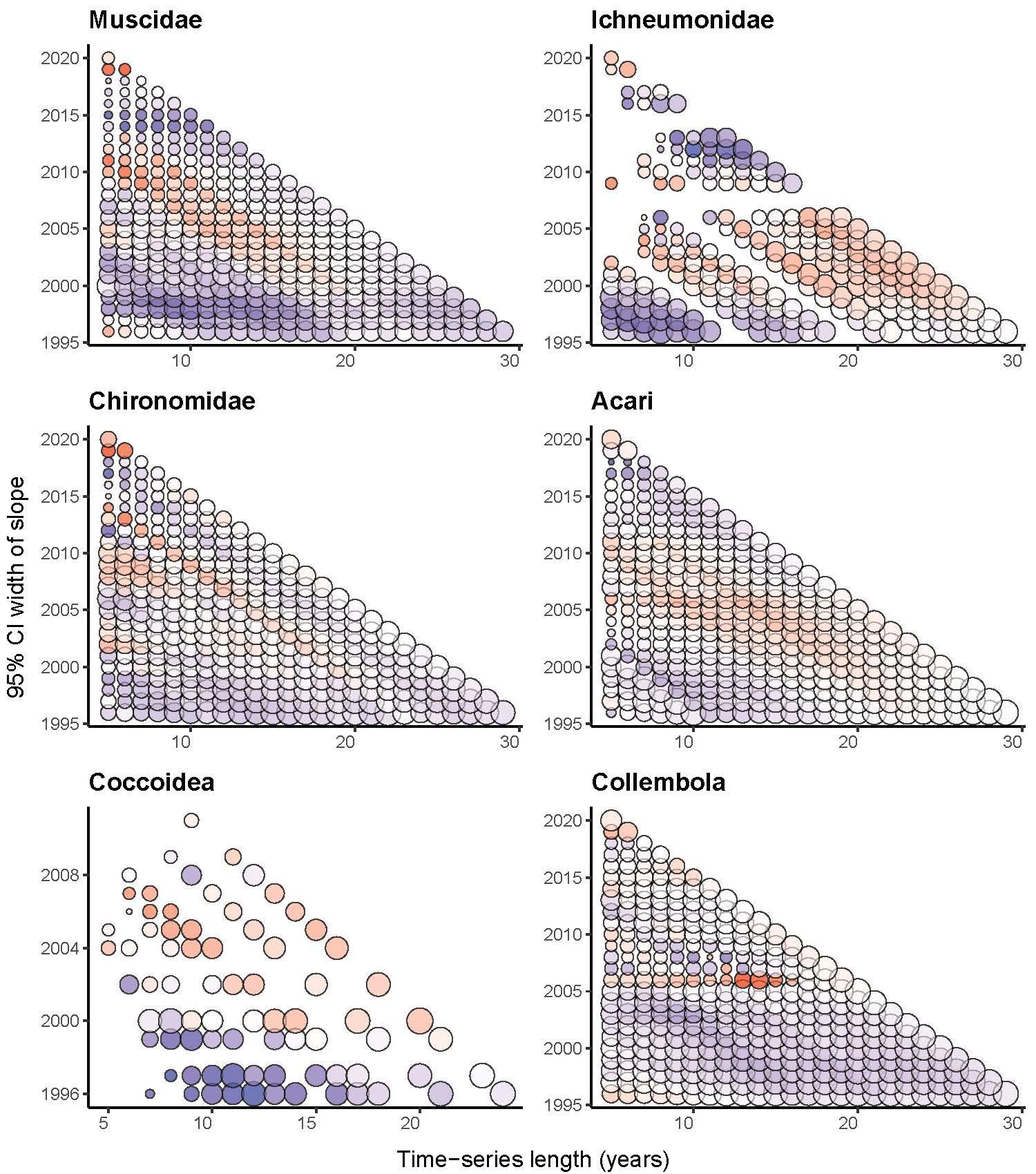

Supplementary Fig. 2. Temporal trends observed at various time-windows of arthropod taxa. The minimum time window was 5 years. Each bubble represents a slope estimated from a sequential model fit. Variation in bubble color and shading indicates directionality and strength of signal of the trend. Darker blue suggests advancing trends while darker red suggests delaying trends. Bubble size indicates the inverse of the 95% credible interval of the slopes, with larger circles representing higher trend certainty. Gaps indicate absence of a slope from a specific window. Representative taxa were selected based on qualitative differences in their apparent phenological patterns to illustrate the range of variation observed across taxa.

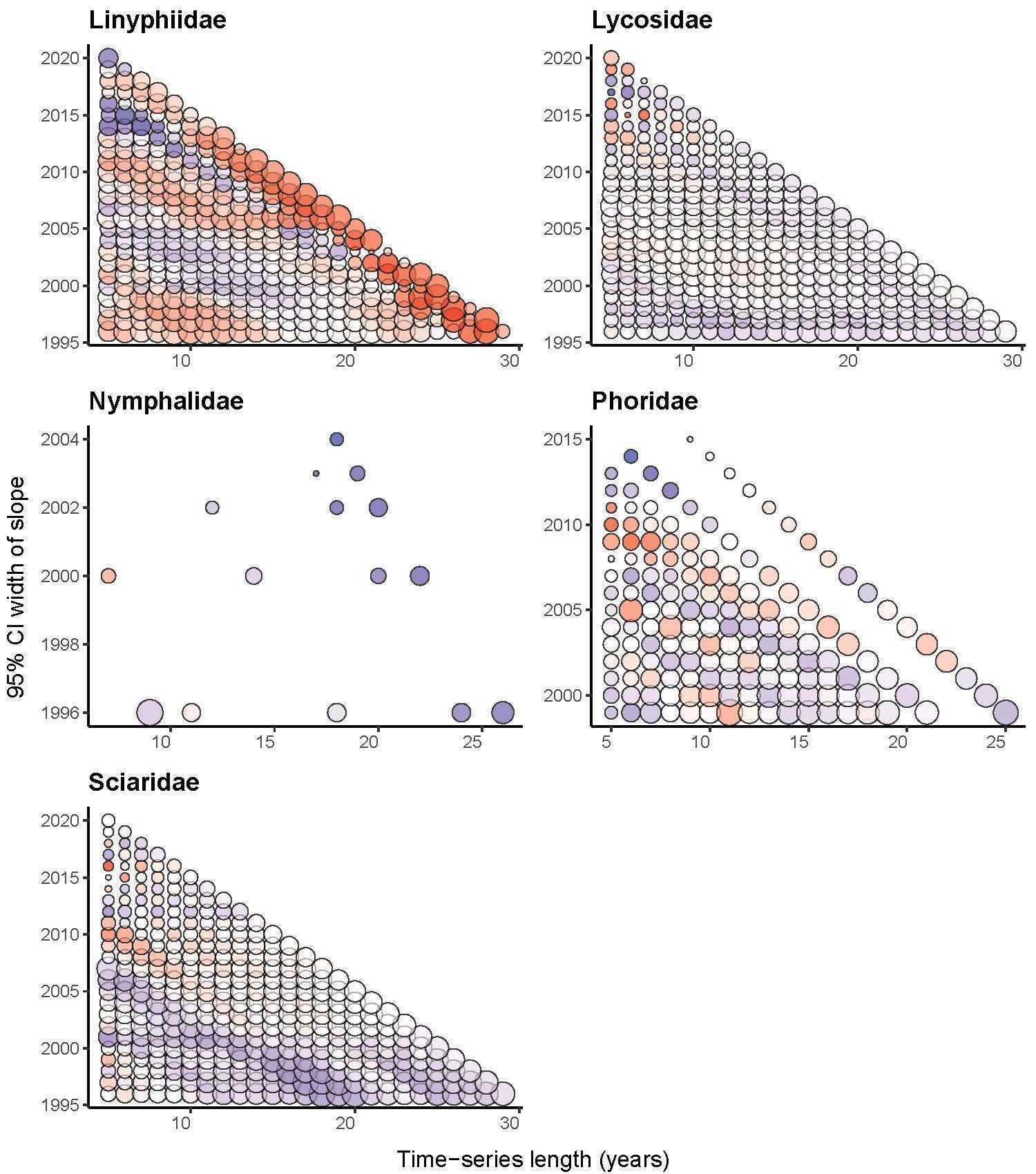

Supplementary Fig. 3. Temporal trends observed at various time-windows of arthropod taxa monitored at Zackenberg Valley. The minimum time window was 5 years. Each bubble represents a slope estimated from a sequential model fit. Variation in bubble color and shading indicates directionality and strength of signal of the trend. Darker blue suggests advancing trends while darker red suggests delaying trends. Bubble size indicates the inverse of the 95% credible interval of the slopes, with larger circles representing higher trend certainty. Gaps indicate absence of a slope from a specific window. Representative taxa were selected based on qualitative differences in their apparent phenological patterns to illustrate the range of variation observed across taxa.

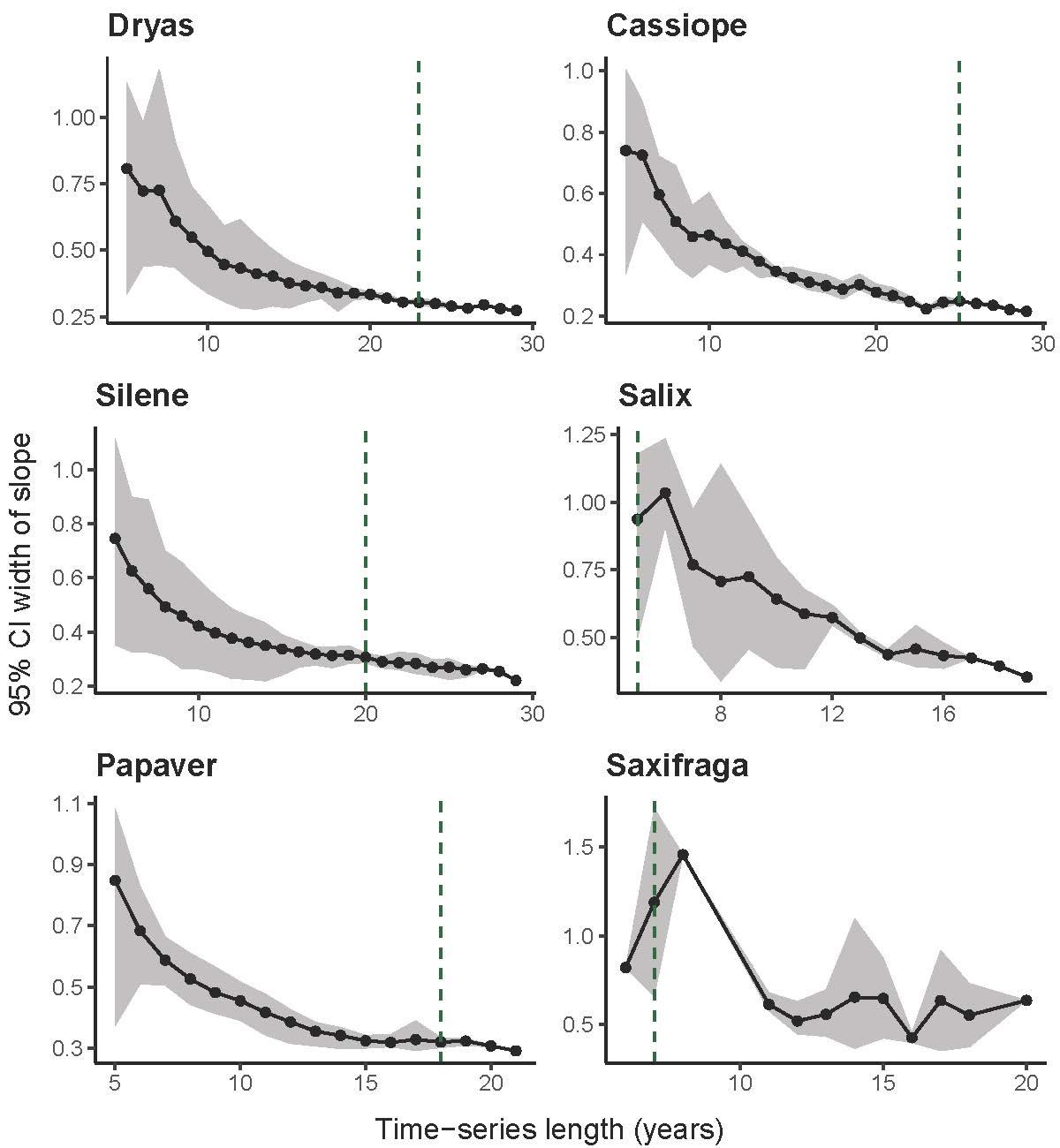

Supplementary Fig. 4. Trend uncertainty over time for all plant taxa. Each point indicates the mean difference between the upper and lower 90% bounds of the slope estimated for a given time-series length, while the gray shading around the line indicates the 95% CI. Dashed lines indicate the minimum time-series length required to recover the direction of long-term phenological change (estimated from full dataset).

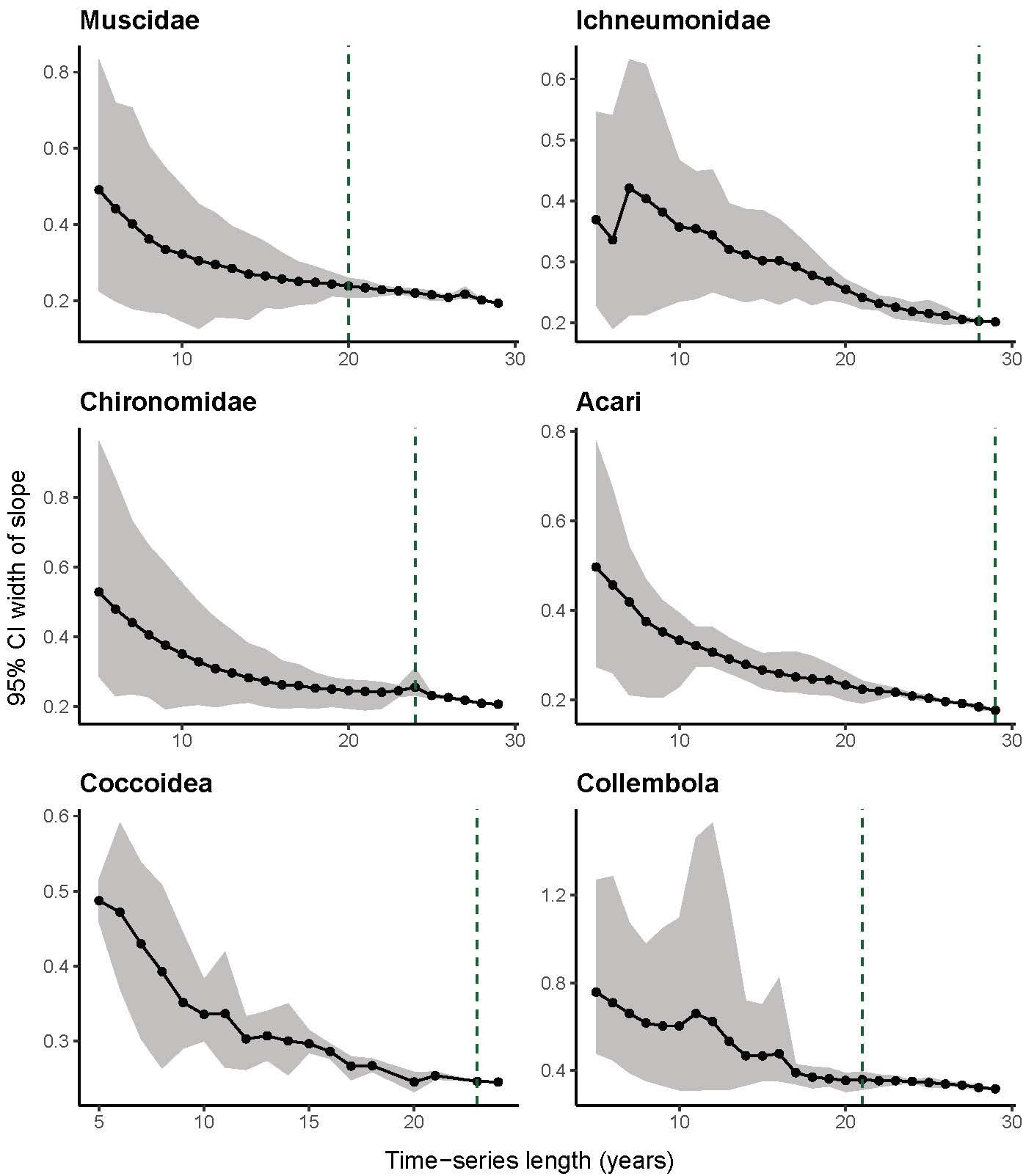

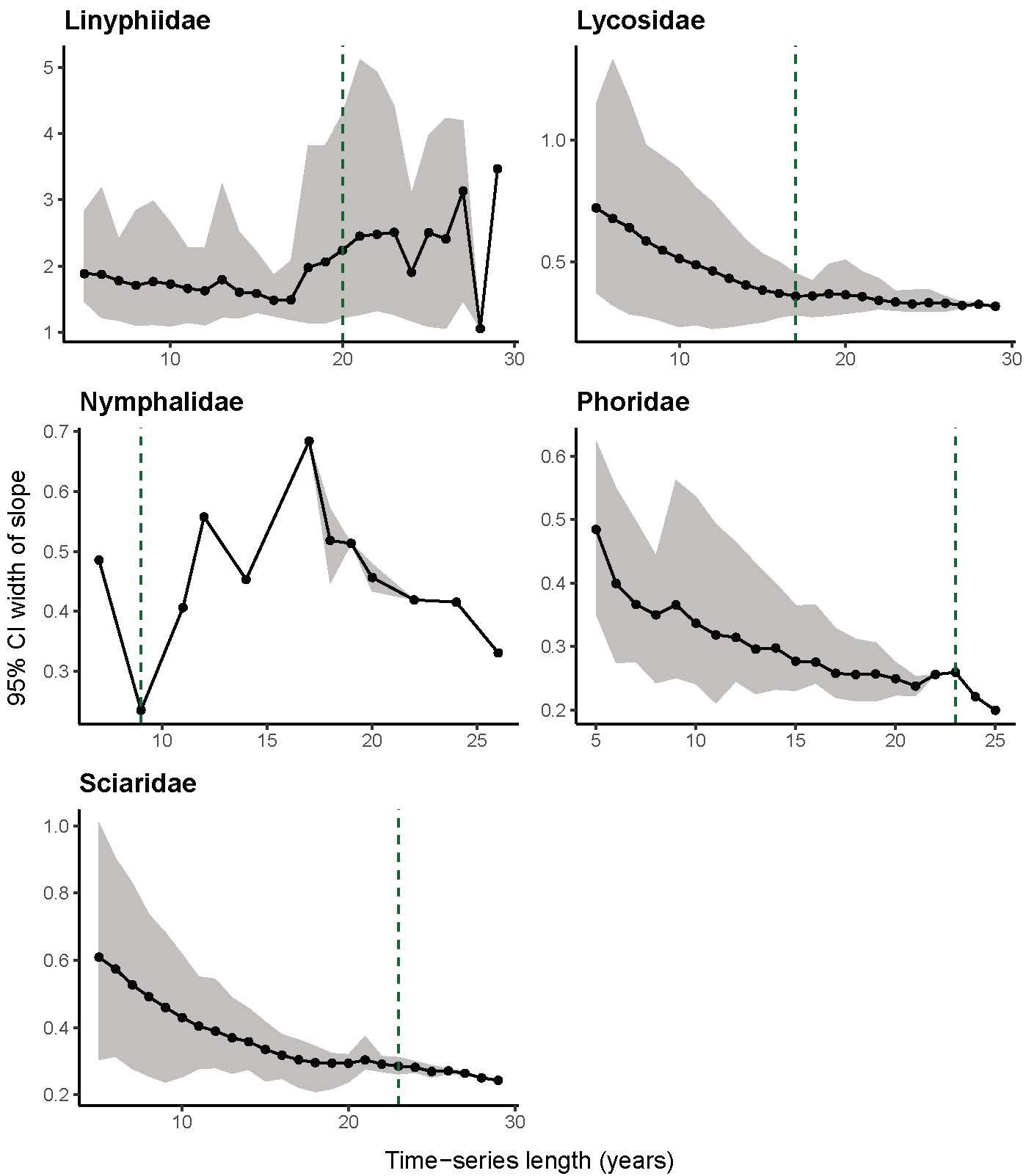

Supplementary Fig. 5. Trend uncertainty over time for all arthropod taxa. Each point indicates the mean difference between the upper and lower 90% bounds of the slope estimated for a given time-series length, while the gray shading around the line indicates the 95% CI. Dashed lines indicate the minimum time-series length required to recover the direction of long-term phenological change (estimated from full dataset).
